# An emergent clade of SARS-CoV-2 linked to returned travellers from Iran

**DOI:** 10.1101/2020.03.15.992818

**Authors:** John-Sebastian Eden, Rebecca Rockett, Ian Carter, Hossinur Rahman, Joep de Ligt, James Hadfield, Matthew Storey, Xiaoyun Ren, Rachel Tulloch, Kerri Basile, Jessica Wells, Roy Byun, Nicky Gilroy, Matthew V O’Sullivan, Vitali Sintchenko, Sharon C Chen, Susan Maddocks, Tania C Sorrell, Edward C Holmes, Dominic E Dwyer, Jen Kok, for the 2019-nCoV Study Group

## Abstract

The SARS-CoV-2 epidemic has rapidly spread outside China with major outbreaks occurring in Italy, South Korea and Iran. Phylogenetic analyses of whole genome sequencing data identified a distinct SARS-CoV-2 clade linked to travellers returning from Iran to Australia and New Zealand. This study highlights potential viral diversity driving the epidemic in Iran, and underscores the power of rapid genome sequencing and public data sharing to improve the detection and management of emerging infectious diseases.

## MAIN TEXT

From a public health perspective, the real-time whole genome sequencing (WGS) of emerging viruses enables the informed development and design of molecular diagnostic methods, and tracing patterns of spread across multiple epidemiological scales (i.e. genomic epidemiology). However, WGS capacities and data sharing policies vary in different countries and jurisdictions, leading to potential sampling bias due to delayed or underrepresented sequencing data from some areas with substantial SARS-CoV-2 activity. Herein, we show that the genomic analyses of SARS-CoV-2 strains from Australian returned travellers with COVID-19 disease may provide important insights into viral diversity present in regions currently lacking genomic data.

### SARS-CoV-2 emergence and dissemination

In late December 2019, a cluster of cases of pneumonia of unknown aetiology in Wuhan city, Hubei province, China was reported by health authorities [1]. A novel betacoronavirus, designated SARS-CoV-2, was identified as the causative agent [2] of the disease now known as COVID-19, with substantial human-to-human transmission [3]. To contain a growing epidemic, Chinese authorities implemented strict quarantine measures in Wuhan and surrounding areas in Hubei province. Significant delays in the global spread of the virus were achieved, but despite these measures, cases were exported to other countries. As of 9 March 2020, these numbered more than 100 countries, on all continents except Antarctica; the total number of confirmed infections exceeded 110,000 and there were nearly 4,000 deaths [4]. Although the vast majority of cases have occurred in China, major outbreaks have also been reported in Italy, South Korea and Iran [5]. Importantly, there is widespread local transmission in multiple countries outside China following independent importations of infection from visitors and returned travellers.

### Whole genome sequencing of SARS-CoV-2 cases in Australia and New Zealand

In New South Wales (NSW), Australia, WGS for SARS-CoV-2 was developed based on an existing amplicon-based Illumina sequencing approach [6]. Viral extracts were prepared from respiratory tract samples where SARS-CoV-2 was detected by RT-PCR using World Health Organization recommended primers and probes targeting the E and RdRp genes, and then reverse transcribed using SSIV VILO cDNA master mix. The viral cDNA was used as input for multiple overlapping PCR reactions (∼2.5kb each) spanning the viral genome using Platinum SuperFi master mix (primers provided in Supplementary Table S1). Amplicons were pooled equally, purified and quantified. Nextera XT libraries were prepared and sequencing was performed with multiplexing on an Illumina iSeq (300 cycle flow cell). In New Zealand, the ARTIC network protocol was used for WGS [7]. In short, 400bp tiling amplicons designed with Primal Scheme [8] were used to amplify viral cDNA prepared with SuperScript III. A sequence library was then constructed using the Oxford NanoPore ligation sequencing kit and sequenced on a R9.4.1 MinION flow-cell. Near-complete viral genomes were then assembled *de novo* in Geneious Prime 2020.0.5 or through reference mapping with RAMPART V1.0.6 [9] using the ARTIC network nCoV-2019 novel coronavirus bioinformatics protocol [10]. In total, 13 SARS-CoV-2 genomes were sequenced from cases in NSW diagnosed between 24 January and 3 March 2020, as well as a single genome from the first patient in Auckland, New Zealand sampled on 27 February 2020 (Table 1). Australian and New Zealand sequences were aligned to global reference strains sourced from GISAID with MAFFT [11] and then compared phylogenetically using a maximum likelihood approach [12].

**Table 1–.** SARS-CoV-2 genomes sequenced in this study.

| **GISAID ID** | **Virus name** | **Location** | **Collection date** | **Travel history** |
| --- | --- | --- | --- | --- |
| EPI_ISL_408976 | 408976/Australia/Sydney-2/2020-01-22 | Sydney, Australia | 22-Jan-20 | China |
| EPI_ISL_407893 | 407893/Australia/NSW01/2020-01-24 | Sydney, Australia | 24-Jan-20 | China |
| EPI_ISL_408977 | 408977/Australia/Sydney-3/2020-01-25 | Sydney, Australia | 25-Jan-20 | China |
| EPI_ISL_413490 | 413490/New_Zealand/01/2020-02-27 | Auckland, New Zealand | 27-Feb-20 | Iran |
| EPI_ISL_412975 | 412975/Australia/NSW05/2020-02-28 | Sydney, Australia | 28-Feb-20 | Iran |
| EPI_ISL_413594 | 413594/Australia/NSW08/2020-02-28 | Sydney, Australia | 28-Feb-20 | SE Asia |
| EPI_ISL_413595 | 413595/Australia/NSW09/2020-02-28 | Sydney, Australia | 28-Feb-20 | SE Asia |
| EPI_ISL_413213 | 413213/Australia/NSW06/2020-02-29 | Sydney, Australia | 29-Feb-20 | Iran |
| EPI_ISL_413214 | 413214/Australia/NSW07/2020-02-29 | Sydney, Australia | 29-Feb-20 | None |
| EPI_ISL_413596 | 413596/Australia/NSW10/2020-02-28 | Sydney, Australia | 01-Mar-20 | SE Asia |
| EPI_ISL_413597 | 413597/AUS/NSW11/2020-03-02 | Sydney, Australia | 02-Mar-20 | Iran |
| EPI_ISL_413600 | 413600/AUS/NSW14/2020-03-03 | Sydney, Australia | 03-Mar-20 | None |
| EPI_ISL_413598 | 413598/AUS/NSW12/2020-03-04 | Sydney, Australia | 04-Mar-20 | Iran |
| EPI_ISL_413599 | 413599/AUS/NSW13/2020-03-04 | Sydney, Australia | 04-Mar-20 | Iran |

### A distinct clade of SARS-CoV-2 identified in travellers returned from Iran

The Australian strains of SARS-CoV-2 were dispersed across the global SARS-CoV-2 phylogeny (Figure 1A). The first four cases of COVID-19 disease in NSW occurred between 24 and 26 January 2020, and these were closely related (with 1-2 SNPs difference) to the prototype strain MN908947/SARS-CoV-2/Wuhan-Hu-1, which is the dominant variant circulating in Wuhan. As the four patients identified in this period had recently returned from China, this region was the likely source of infection. From 1 February 2020, travel to Australia from mainland China was restricted to returning Australian residents and their children, who were placed in home quarantine for 14 days. Despite the intensive testing of such returning travellers, no further cases of COVID-19 were detected in NSW until 28 February 2020, when SARS-COV-2 was detected in an individual returning from Iran (NSW05). A close contact of this individual also tested positive (NSW14) providing the first evidence of local transmission within NSW. This was followed by further Iran travel-linked cases in NSW (NSW06, NSW11, NSW12, NSW13) and New Zealand (NZ01).

**Figure 1.**
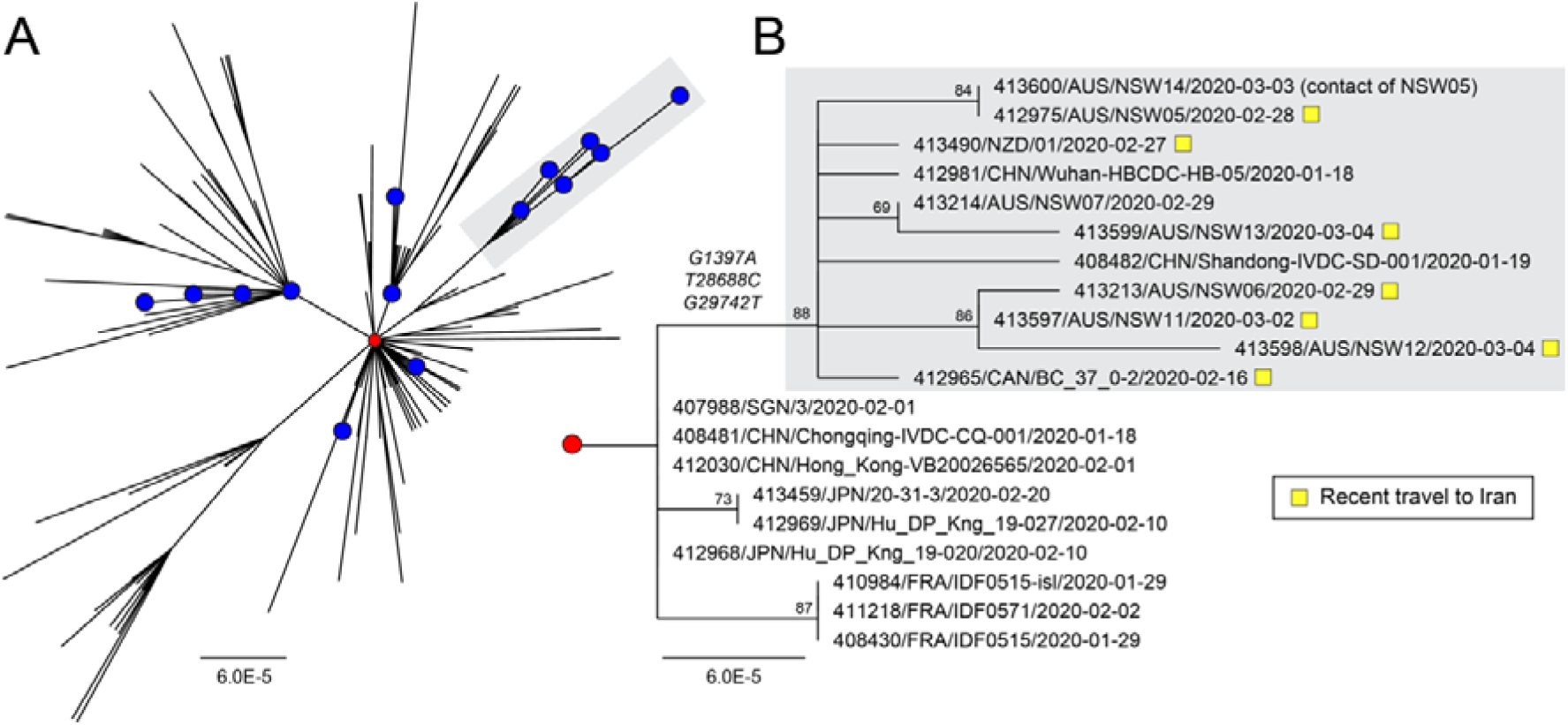
Phylogenetic analysis of SARS-CoV-2 genome sequences highlighting a clade of imported cases from Iran. (A) Global diversity of circulating SARS-CoV-2 strains including Australian sequences (blue circles, n=19). The prototype strain Wuhan-Hu-1 is shown as a red circle. An emergent clade containing cases imported from Iran is highlighted with grey shading. (B) Sub-tree showing the informative branch containing imported Iranian cases (highlighted with yellow squares) and defined by substitutions at positions G1397A, T28688C, G29742T. Node support is provided as bootstrap values of 100 replicates. For both panels A & B, the scales are proportional to the number of substitutions per site.

Of note, the genomes of all patients with a history of travel to Iran were part of a monophyletic group defined by three nucleotide substitutions (G1397A, T28688C & G29742T) in the SARS-CoV-2 genome relative to the Wuhan prototype strain (Figure 1B). G1397A and T28688C both occur in coding regions with G1397A producing a non-synonymous change (V378I) in the ORF1ab encoded non-structural protein 2 region. G29742T occurs in the 3’ UTR. In addition to the Australian and New Zealand strains, this clade also included a traveller who had returned to Canada from Iran (BC_37_0-2), providing further evidence of its likely link to the Iranian epidemic. Indeed, a search of all currently available GISAID sequences and metadata revealed no other complete genome sequences from patients with documented history of travel to or residence in Iran (as of 9 March 2020). A search of partial sequences identified two SARS-CoV-2 sequences which originated in Iran (413553/IRN/Tehran15AW/2020-02-28 and 413554/IRN/Tehran9BE/2020-02-23) spanning a 363 nt region of the viral nucleoprotein (N). Although short in length, these two sequences covered one of the informative SNPs defining this clade - T28688C, and both Iranian strains matched the sequences from patients with travel histories to Iran and grouped by phylogenetic analysis (Supplementary Figures S1 & S2).

## Discussion

Technological advancements and the wide-spread adoption of WGS in pathogen genomics have transformed public health and infectious disease outbreak responses [13]. Previously, disease investigations often relied on the targeted sequencing of a small locus to identify genotypes and infer patterns of spread along with epidemiological data. As seen with the recent West African Ebola [14] and Zika virus epidemics [15], rapid WGS significantly increases resolution of diagnosis and surveillance thereby strengthening links between clinical and epidemiological data [16]. This advance improves our understanding of pathogen origins and spread that ultimately lead to stronger and more timely intervention and control measures [17]. Following the first release of the SARS-CoV-2 genome [18], public health and research laboratories worldwide have rapidly shared sequences on public data repositories such as GISAID [19] (n = 236 genomes as of 9 March 2020) that have been used to provide near real-time snapshots of global diversity through public analytic and visualization tools [20].

While all known cases linked to Iran are contained in this clade, it is important to note the presence of two Chinese strains sampled during mid-January 2020 from Hubei and Shandong provinces. It is expected that further Chinese strains would be identified within this clade, and across the entire diversity of SARS-CoV-2 as this is where the outbreak started, including for the outbreak in Iran itself. However, while we cannot completely discount that the cases in Australia and New Zealand came from other sources including China, our phylogenetic analyses, as well as epidemiological (recent travel to Iran) and clinical data (date of symptom onset), provide evidence that this clade of SARS-CoV-2 is linked to the Iranian epidemic, from where genomic data is currently lacking. Importantly, the seemingly multiple importations of very closely related viruses from Iran into Australia suggests that this diversity reflects the early stages of SARS-CoV-2 transmission within Iran.

## ACKNOWLEDGMENTS

The members of the nCoV-2019 Study Group include Linda Donovan, Shanil Kumar, Tyna Tran, Danny Ko, Christine Ngo, Tharshini Sivaruban, Verlaine Timms, Connie Lam, Mailie Gall, Karen-Ann Gray, Rosemarie Sadsad and Alicia Arnott. The authors acknowledge the Sydney Informatics Hub and the use of the University of Sydney’s high performance computing cluster, Artemis, and all the laboratories that have referred SARS-CoV-2 samples to the Centre for Infectious Diseases and Microbiology Laboratory Services, NSW Health Pathology - Institute of Clinical Pathology and Medical Research, Westmead Hospital.

We would finally like to thank all the authors who have kindly shared genome data on GISAID, and have included a table (Supplementary Table S2) outlining the authors and institutes involved. Data including the sequences in this study are available for download from https://www.gisaid.org/.

## AUTHOR CONTRIBUTIONS

Study concept and design by JSE, ECH & JK. Sample processing and testing by IC & HR. Sequencing and analysis by JSE, RR, JDL, JH, MS, XR, RT & ECH. Study coordination by NG, MVOS, VS, SCC, SM, TCS, DED & JK. JSE wrote the first manuscript draft with editing from ECH, JDL, RR, TCS, VS & JK. The final manuscript was approved by all authors.

